# Minimally invasive deep-brain imaging through a 50 μm-core multimode fibre

**DOI:** 10.1101/289793

**Authors:** Sebastian A. Vasquez-Lopez, Vadim Koren, Martin Plöschner, Zahid Padamsey, Tomáš Čižmár, Nigel J. Emptage

## Abstract

Achieving optical access to deep-brain structures represents an important step towards the goal of understanding the mammalian central nervous system. The complex refractive index distribution within brain tissue introduces severe aberrations to long-distance light propagation thereby prohibiting image reconstruction using currently available non-invasive techniques. In an attempt to overcome this challenge endoscopic approaches have been adopted, principally in the form of fibre bundles or GRIN-lens based endoscopes. Unfortunately, these approaches create substantial mechanical lesions of the tissue precipitating neuropathological responses that include inflammation and gliosis. Together, lesions and the associated neuropathology may compromise neural circuit performance. By replacing Fourier-based image relay with a holographic approach, we have been able to reduce the volume of tissue lesion by more than 100-fold, while preserving diffraction-limited imaging performance. Here we demonstrate high-resolution fluorescence imaging of neuronal structures, dendrites and synaptic specialisations, in deep-brain regions of living mice. These results represent a major breakthrough in the compromise between high-resolution imaging and tissue damage, heralding new possibilities for deep-brain imaging *in vivo*.

## Introduction

At present, non-invasive (surface) high-resolution imaging of brain tissue can achieve micrometre resolution up to penetration depths of about 1 mm^1^. Even in small mammals, such a mice, this severely restricts optical access to almost all subcortical structures many of which are implicated in important neuronal processes such as memory formation and gating of sensory and motor information, as well as neurological diseases, for example Alzheimer’s disease^2–4^. Attempts to gain access to these brain regions have precipitated the use of invasive strategies, including the removal of overlying cortical structures^5^, the insertion of fibre bundles^6^ and graded index (GRIN) lenses^7,8^. However, where these techniques are uniformly problematic is the considerable extent to which they damage the brain^9^. The degree to which this damage is consequential for the behaviour of the animal and the physiology of networks is not fully understood^10,11^, nonetheless, it is clear that were it possible to cause little or no damage, then such an approach would be wholly desirable as a method with which to explore the nervous system *in vivo*.

Advances in holographic methods and computational power have recently allowed for a novel approach to high-resolution imaging, utilising deterministic light propagation through optically complex media and, of particular importance for this work, multimode optical fibres (MMF)^12–14^. MMF probes offer the advantage that they are many times thinner than other types of microendoscopes. To exploit this advantage we have developed a compact and highly optimised approach for minimally invasive *in vivo* brain imaging applications, utilising holographic control of light propagation through a single 50 μm-core multimode fibre. In this paper we present the first demonstration of this technique applied *in vivo*, by acquiring fluorescent images of neuronal structures from regions located deep in the living rodent brain.

## Results

The principles behind the MMF imaging method are detailed in previous reports^12,13^. Integral to the system is a computer-controlled dynamic optical element, a spatial light modulator (SLM), which enables us to manipulate the propagation of light at will, through the whole optical train comprising an arbitrary length of MMF. Prior to commencement of imaging, the SLM is used in a calibration procedure during which we acquire a transmission matrix (TM) of the optical system^15^. The availability of the system-specific TM then allows us to produce a set of holographic modulations, which are employed in the image acquisition procedure. Each of these modulations, when applied at the SLM, produces a diffraction-limited spot at a specific location across the fibre output plane. Importantly, spots may be generated at an arbitrary distance from the distal fibre facet. This combination of procedures provides the basis for fibre-based volumetric scanning fluorescence microendoscopy, achieving diffraction-limited performance with an endoscopic probe whose diameter is several fold smaller than any used previously. This represents an enormous step in the trade-off between image resolution and device footprint (Fig. 1b, c).

**Figure 1.**
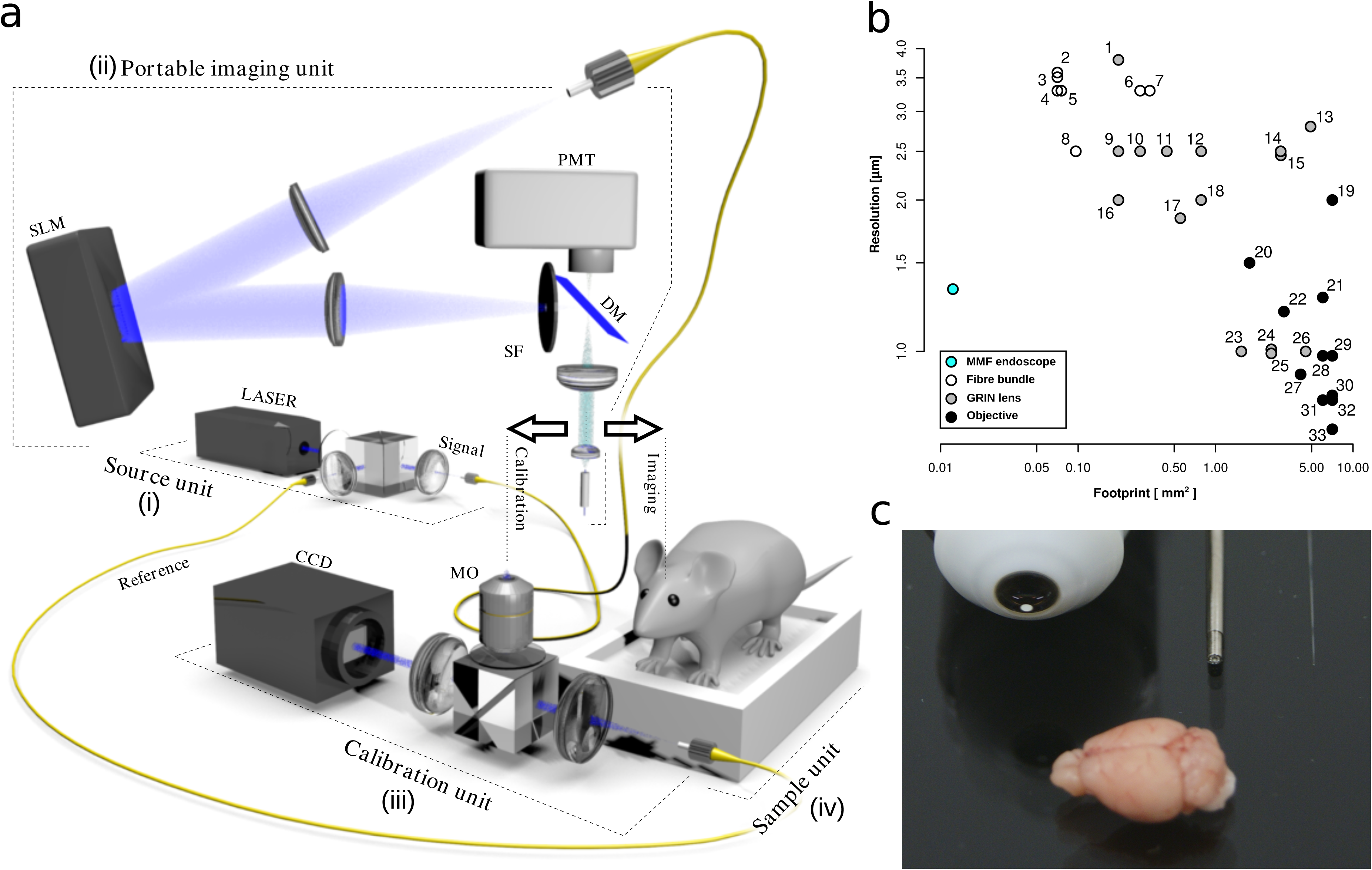
A multimode fibre (MMF) imaging system for minimally invasive deep-brain *in vivo* imaging. **a**, Schematic diagram of the experimental geometry. (i) Laser power distribution unit for excitation at 532 nm or 488 nm. (ii) The main portable imaging arm controls the light propagation through the MMF by means of phase manipulation of light at individual pixels of a spatial light modulator (SLM). The calibration unit is used for the acquisition of the transmission matrix prior to imaging. The sample unit holds tissue sections and head fixed animal models. *DM*, dichroic mirror; *MMF*, multimode fibre; *MO*, microscope objective; *PMF*, polarization maintaining fibre; *PMT*, photo multiplier tube; *SF*, single-mode fibre. **b**, The relationship between image resolution and instrument footprint for *in vivo* light-based imaging modalities. The MMF data point is shown in blue. Data for other imaging modalities are taken from the references listed in Supplementary Table 1. **c**, A direct comparison of the size of a mouse brain with *in vivo* imaging devices: a x60 water immersion objective, a GRIN lens and the 50µm core MMF used in this study.

The optical geometry has been optimised to provide the functional stability and mobility necessary for use *in vivo* (Fig. 1a). The initial section of the system distributes laser light of 532 nm or 488 nm into two single-mode polarisation-maintaining optical fibres, one to deliver the excitation signal to the SLM and the other to provide a reference signal during calibration. The main optical arm, comprising the SLM, the MMF probe, relay optics and a fluorescence detection unit, has been designed to be compact and is embedded within a robust caged framework housed on a 3-dimensional micro-positioning stage to facilitate alignment during calibration as well as navigation of the MMF probe into the brain tissue. Finally, the calibration arm, used only during TM acquisition, relays the MMF probe output signal at the CCD and combines this with the reference beam. The implementation of a GPU-accelerated toolbox for SLM control^14^, allows us to acquire the full transformation matrix of a 50 μm-core MMF in less than 4 minutes. Imaging can be performed immediately after the TM acquisition and, as long as the fibre alignment is not changed, continued for hours and even days without the need for further calibration.

To assess the spatial resolution of the imaging system, we show images of a custom-made USAF-1951 resolution target (Fig. 2a). Figure 2b shows that at 532 nm the spatial features can be resolved down to 1.5μm, which agrees well with the Rayleigh criterion for imaging systems with corresponding numerical aperture of the fibre (0.22). Migration to a wavelength of 488 nm enhances the resolution proportionally, ≈1.35μm. Of key relevance to our primary objective, accurate imaging of neuronal structures, we wish to stress that our scanning approach is free from the granulous artefacts commonly observed when using computational approaches^16,17^, as these can be easily confused with synaptic structures. This is a particular concern where high-resolution images are collected from sparsely labelled samples over a small area, as it is frequently used in neurobiology^18,19^. Increasing the content of light emitting structures across the field of view in our approach may intensify the magnitude of a uniform background signal, however, the resolution is fully maintained while introducing no artefacts to imaged structures (Fig. 2a). In order to assess the performance of our system when imaging neuronal structures, we first conducted imaging trials in *ex vivo* brain slices from the rat hippocampus (Fig. 2c,d,e). Fluorescently labelled neurons were imaged with both a standard confocal microscope equipped with a 60× water immersion objective (Fig. 2d,e green images) and our fibre-based system (Fig. 2d,e grey images). Dendritic spines and axonal boutons are clearly visible, with identical structures identifiable in both MMF and confocal images, suggesting that the device would be suitable for structural imaging studies^19^. This offers robust validation of the MMF approach for the acquisition of fluorescent images in living tissue. Furthermore, even though the fluorescence yield varies significantly between the axonal and dendritic arbour, adjustments in sensitivity of the apparatus allow for signal recording without major loss of resolution.

**Figure 2.**
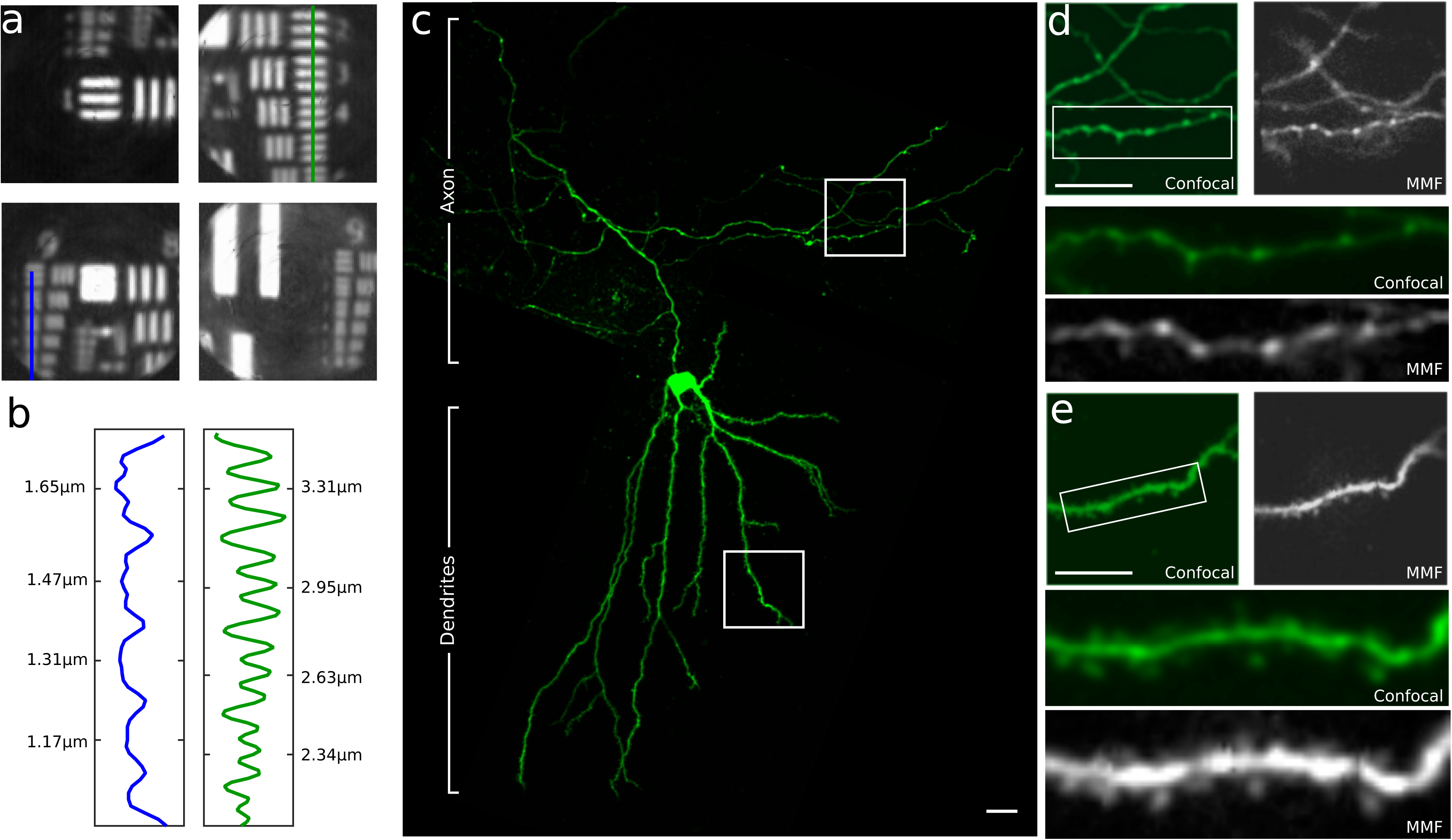
Resolution measurements and comparison of MMF and confocal imaging in *ex vivo* tissue of the hippocampus. **a-b**, Resolution measurements were conducted on a custom USAF-1951 resolution target. The resulting lateral resolution of ≈ 1.5 μm has been confirmed at the wavelength of 532 nm. **c**, Confocal imaging of a hippocampal neuron in a brain slice was collected using a Biorad confocal microscope equipped with a 60× water immersion objective. **d-e**, Structural images were obtained through the MMF (d,e grey images) and the confocal microscope (d,e green images) of same regions of the neuron to facilitate direct comparison. Axonal boutons (d) and dendritic spines (e) can be clearly identified using each imaging modality, with little evidence of data loss for the MMF imaging. Scale bars = 20 µm.

Having demonstrated that the system performs well when imaging living neuronal tissue *ex vivo*, we set out to explore whether we could achieve comparable results *in vivo.* Here we sought to image neurons from deep regions of the intact brain of live mice. We used the Thy1-GFP-M mouse transgenic line that expresses fluorescently labelled neurons sparsely throughout the nervous system; an approach commonly used for *in vivo* neuronal structural imaging studies^18^. As the diameter of our fibre is small, 125 μm inclusive of cladding, we were able to insert the fibre directly into the brain tissue via a small craniotomy and image ‘live’ as we advanced slowly through the tissue. This is a significant improvement over existing methods for deep-brain imaging that require extensive surgery and the aspiration of overlying brain tissue. On identifying our target structure we were able to begin imaging immediately, again a marked contrast with other endoscopic brain imaging methods that require several days post-surgery before imaging can commence^9^. Critically, we see little evidence of damage to blood vessels and so images were not obscured by tissue bleeding.

Figure 3 shows images of a fluorescent neuron in the dorsal striatum imaged after lowering a fibre 1.5 mm into the brain of an anaesthetised mouse and collecting an image stack at different focal planes beneath the fibre (Fig. 3a,b). A dendritic branch on which there are dendritic spines is clearly identifiable (Fig. 3c) with the three-dimensional structure becoming readily apparent with holographic progression of the focal plane. The ability to adjust the focal plane while maintaining the fibre at a fixed position represents a further considerable advantage of the MMF fibre system for *in vivo* imaging. The imaging plane can be adjusted over a range 0-100 μm from the fibre facet with no movement of the fibre and therefore no mechanical consequence for the brain tissue. This further minimises the impact of fibre placement into the tissue. The extent to which the placement of the fibre impacts upon the neuronal tissue during image collection is shown in a post mortem section of brain tissue (Fig. 3d). Labelled neurons and their dendritic processes remain alive and intact even when located in close proximity to the fibre tract (Fig 3d inset).

**Figure 3.**
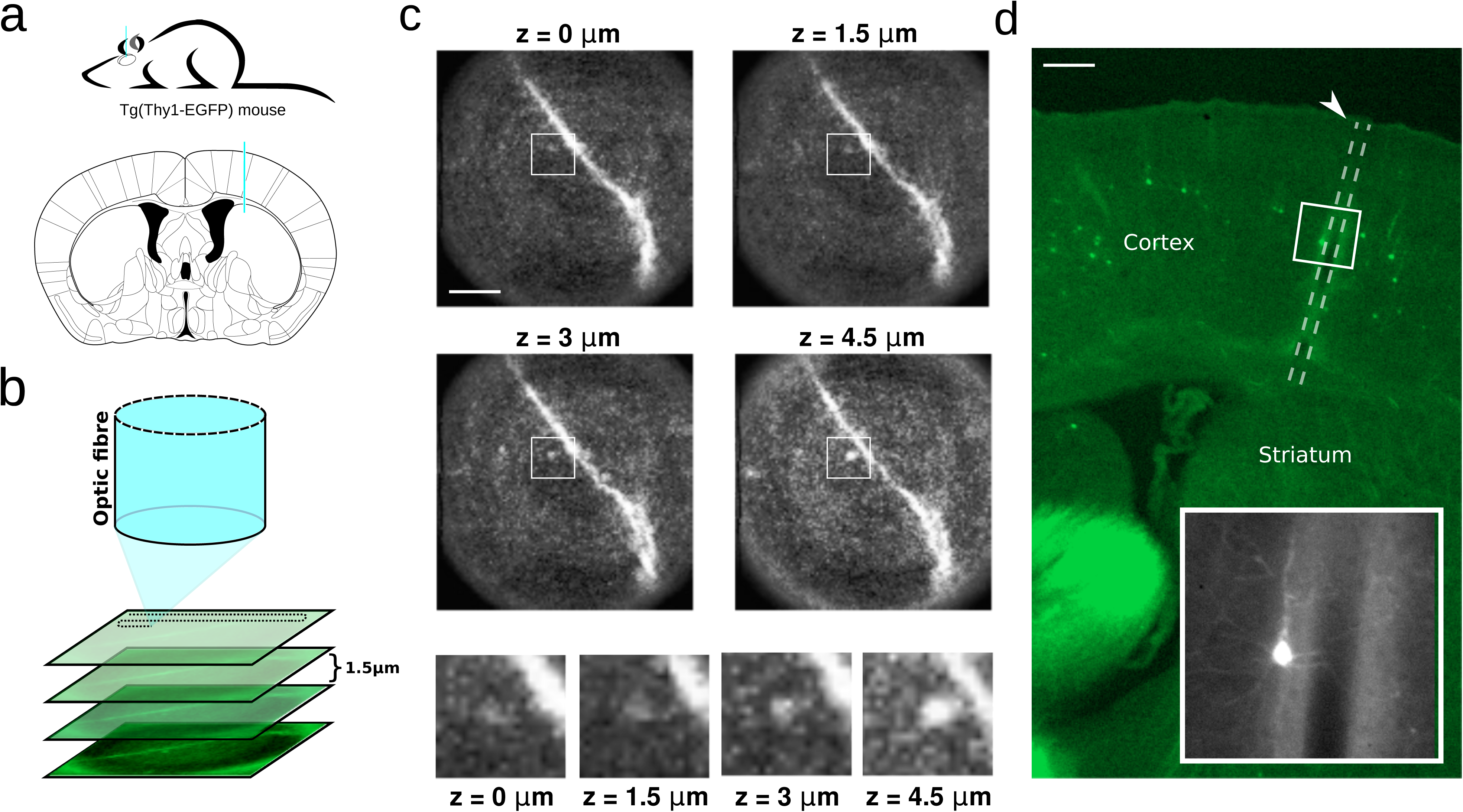
Fibre-based structural imaging of a GFP-labelled dendrite of a neuron in the dorsal striatum of an anaesthetised mouse. **a**, Imaging was performed by lowering a multi-mode fibre 1.8 mm into the brain of an anaesthetised mouse (top). Atlas depiction of the region of the striatum imaged in (c) adapted from the Allen Mouse Brain Atlas (bottom) with fibre placement in blue. **b**, Fluorescence imaging can be performed at multiple depths from the front facet of the fibre by calibrating the system to different focal planes. **c**, Dendritic spines are clearly identifiable and their three-dimensional structure becomes readily apparent by varying the focal plane. Scale bar = 10 µm. **d**, Post mortem histological sectioning of the mouse brain imaged in (c) shows the path of the fibre (arrow) through the cortex, with minimal damage to the tissue. Scale bar = 200 µm. Image inset shows that the structure of cortical neurons is preserved even around the margins of the fibre track.

While structural imaging is a valuable technique, we wished to assess whether fibre-based dynamic imaging can also be achieved. Figure 4a shows that our system can detect slow changes in fluorescence elicited in a neuron loaded with the Ca^2+^ indicator OGB-1, in *ex vivo* brain slices from the rat hippocampus. For *in vivo* imaging, neurons of the medial geniculate body (MGB)—a part of the auditory thalamus— of C57BL/6 mice were sparsely labelled with the Ca^2+^ reporter GCaMP6m, in order to record action potentials elicited in response to auditory stimuli. Mindful that the scan rate of the system would not allow for full-frame fast functional imaging (frame rate = 2.4 seconds per 120×120-pixel frame) in the current configuration, we instead imaged a small group of pixels at 33.3 Hz (chosen to reflect the scan rate of commercial 2-photon systems). Using this approach, we were able to observe robust and reproducible stimulus-driven Ca^2+^ signals from sound-responsive neurons in anesthetized mice *in vivo* (Fig 4b,c,d).

**Figure 4.**
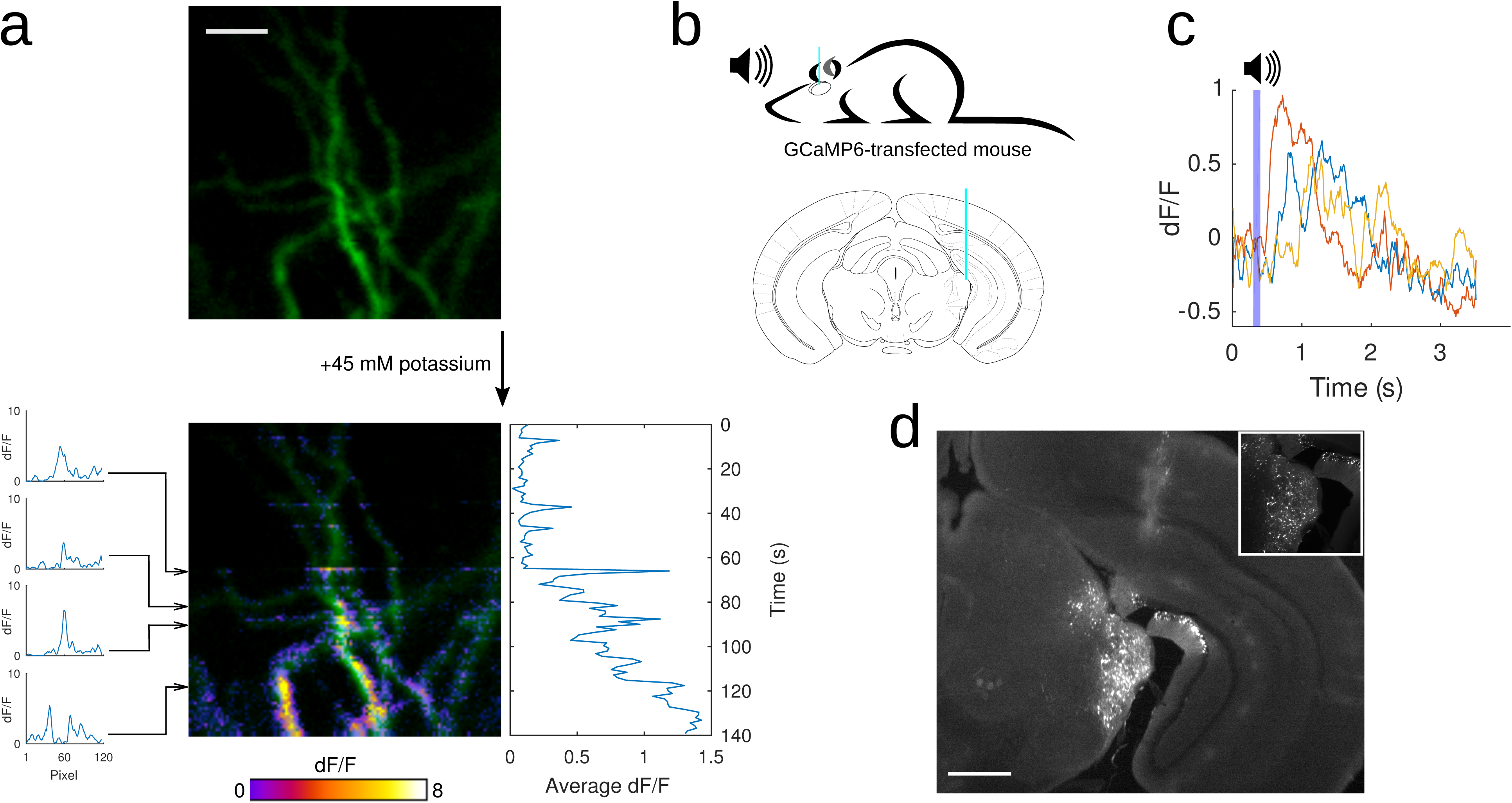
Fibre-based imaging of dynamic changes in neuronal Ca^2+^ signals. **a**, A neuron in an organotypic hippocampal slice was filled with the Ca^2+^-sensitive dye OGB-1 (top). After bath application of 45 mM of potassium the gradual increase in intracellular Ca^2+^ resulting from the depolarisation of the neuron was detected through dynamic changes in fluorescence (bottom). Scale bar = 10 μm. **b**, A multimode optical fibre was lowered 3 mm into the medial geniculate body (MGB) of an anaesthetised mouse (top). Atlas depiction indicating the placement of the MMF; adapted from the Allen Mouse Brain Atlas (bottom) with fibre placement in blue. **c**, Calcium responses recorded from a single pixel were elicited by repeated presentation of a 100 ms pure tone of 16 kHz (blue bar). Colour traces show responses from individual trials. **d**, Post-mortem histological analysis shows sparse expression of the genetically-encoded calcium indicator GCaMP6m in the MGB and the fibre track. Inset shows the level of sparse labelling of MGB neurons. Scale bar = 1 mm.

## Discussion

We have achieved *in vivo* fluorescent imaging of neurons within deep-brain structures of mice, which we believe to be the least invasive, deep-brain, high-resolution approach reported to date. This method provides a route to achieving high-resolution optical access to deep-brain sub-cellular processes in living and ultimately freely behaving animals with minimal disruption to associated circuitry. This is also a prominent demonstration of applications of wavefront-shaping microscopy in biomedical research^20^ and an overture for future advances in minimally-invasive imaging *in vivo*^21^.

In its current configuration, the imaging system can operate at a scan rate of approximately 10 ms per pixel, determined by the maximum refresh rate of 100 Hz for the SLM. Faster scan systems based around acousto-optic deflectors (AOD)^13^ or digital micromirror devices^22,23^ are available and could be incorporated into our geometry. Here we show early data that the fibre-based system is capable of detecting dynamic changes in fluorescence signals that correlate with neuronal activity *in vivo*. Future developments will now aim to achieve the scanning speeds necessary for wide-field imaging enabling neuronal population activity to be monitored. Enhancement in scan speed will also be essential for imaging in freely moving animals using extensive lengths of fibre. We have previously shown^24^ that the changes in the transmission matrix caused by fibre bending can be corrected if fibre shape is accurately monitored. Of the remaining challenges for *in vivo* functional imaging comes from the need to avoid contamination of the dynamic signal from unwanted epifluorescent sources^25^. Some progress has been made in this regard by implementing a form of holographic confocal microscopy^26^ as well as by demonstrating the potential of step index multimode fibres for achieving fibre-based two-photon microscopy^27^.

## Methods

### Experimental geometry

A monochromatic, linearly polarised light beam (λ = 532/488 nm; CrystaLaser) is split and coupled into two separate optical fibres (a polarisation maintaining single-mode fibre SMF). Light exiting the SMF is collimated (f = 60mm, achromatic doublet) and its polarisation is aligned with working polarisation of the SLM (Meadowlark Optics, 512 × 512 pixels) using a half-wave plate. The light is phase modulated by the SLM in an off-axis regime and Fourier transformed by a plano-convex (f = 100mm) lens onto an iris that only transmits the first diffraction order. The transmitted signal is then reflected by a dichroic mirror (Thorlabs MD498) and circularly polarised by a quarter-wave plate to assure minimum coupling between polarisation states. Circularly polarised light then enters a telescope consisting of plano-convex (f = 50mm) and aspheric (f = 8mm) lenses coupling the light into a multimode fibre (Thorlabs, FG050UGA NA = 0.22). The subsequent light path depends on the mode of operation of the optical system. The standard experimental protocol consists of two modes of operation: acquisition of TM (system calibration), and image acquisition.

### Acquisition of TM

In this mode, the light exiting the multimode fibre is coupled into the calibration unit. Here the light output is imaged by a microscope objective lens (Olympus 20×, NA = 0.4) and an achromatic doublet (f = 150mm) onto a CCD camera (Basler pilot piA640-210gm). In-between the lenses, the signal is converted back into the linear polarisation state using a quarter-wave plate and merged with a reference signal using a 50:50 non-polarising beam-splitter. TM is a linear relation between bases of input and output modes^15^. In our implementation, the input modes are diffraction-limited focal points defined across an orthogonal grid (50×50) at the input facet of the MMF and the basis of 120×120 output modes is analogously defined along the plane of the MMF output facet or any plane axially displaced away from the fibre facet thus allowing for volumetric imaging (sectioning). The SLM is used to sequentially generate input modes that propagate through the MMF and leave as a linear combination of output modes. These undergo further sequential phase offsetting and interference with the reference signal. The output modes are analysed by a corresponding number (120×120) of CCD camera pixels, thus resulting in one vector of the TM matrix. After acquisition of all input modes the TM measurement is completed and can be used to design input fields (and the corresponding SLM modulation) necessary for generation of individual output modes (diffraction-limited foci at the MMF output) or any other desired light field leaving the MMF. Further details of the approach are available elsewhere^14^.

### Point-scanning-based image acquisition

Once calibrated, the microscope objective beneath the MMF is exchanged for the sample unit, this houses *ex vivo* neural tissue or the anaesthetised animal. Once in position the fibre is lowered into the area of interest within the tissue. The SLM’s diffraction grating generates a holographic projection across the proximal fibre facet. This linear superposition of focal points, constructively interferes while propagating through the fibre in order to collimate into a single, diffraction limited focal light-point at short distance from the distal fibre end. With a maximum SLM refresh rate of 204Hz, this point can be used to raster-scan a set of 120×120 output modes or their arbitrary selection across the fluorescent sample, whereby the emitted wavelength is collected and passed back through the fibre. The intensity of the transmitted response signal is registered for each raster scan position by a photo multiplier tube (PMT) and constitutes the pixel value in the final acquired image.

### Ex vivo imaging

All animal work was carried out in accordance with the Animals (Scientific Procedures) Act, 1986 (UK), and under project and personal licenses approved by the Home Office (UK). Organotypic hippocampal brain slices (350 μm) were prepared from male Wistar rats (postnatal day 7). Slices were plated on Millicell inserts (Millipore) and incubated at 34°C and 5% CO2 for 7-14 days prior to use. Slices were maintained in 1mL of culture medium (50% Minimum Essential Media, 25% heat-inactivated horse serum, 23% Earl’s Balanced Salt Solution, and 2% B-27 with 6.5 g/L added glucose; ThermoFisher Scientific), which was replaced every 2-3 days. Slices were imaged at room temperature in 1 mL of physiological Tyrode’s solution (in mM: 120 NaCl, 2.5 KCl, 30 glucose, 2 CaCl_2_, 1 MgCl_2_, and 25 HEPES, with 2 ascorbic acid and 1 Trolox added to minimize photodynamic damage; Sigma Aldrich; pH = 7.2-7.4). Dentate granule neurons were loaded with fluorescent dye using whole-cell patch electrophysiology. Briefly, glass electrodes (4-8 MΩ resistance) were filled with standard internal solution (in mM: 135 KGluconate, 10 KCl, 10 HEPES, 2 MgCl_2_, 2 Na_2_ATP and 0.4 Na_3_GTP; pH = 7.2-7.4) containing either 2 mM Alexa Fluor 488 fluorescent dye (ThermoFisher Scientific) for structural imaging or 1 mM of the Ca2+ sensitive dye, Oregon Green BAPTA-1 (ThermoFisher Scientific), for functional imaging. Cells were patched for 5-10 minutes to give adequate time for dye diffusion, after which the patch electrode was slowly retracted, enabling the plasma membrane to reseal. Structural and functional images were obtained through an optic fibre. During functional imaging, 500 μl of a high potassium Tyrode’s solution (in mM: 32.5 NaCl, 90 KCl, 30 glucose, 2 CaCl_2_, 1 MgCl_2_, and 25 HEPES, with 2 ascorbic acid and 1 Trolox added to minimize photodynamic damage; pH = 7.2-7.4) was added to the bath to drive spontaneous neuronal activity. For purposes of comparison, structural images were additionally acquired using a confocal microscope (BioRad Radiance 2000) equipped with a 488 nm argon laser line, and controlled by LaserSharp (BioRad) software. Confocal images were acquired as a high resolution (1024 × 1024 pixels) z-stack (1 μm step size), taken through a 60x water-immersion objective (0.9 NA; Olympus) on an upright Olympus BX50WI microscope.

### In vivo imaging

All experiments were approved by the local ethical review committee at the University of Oxford and licensed by the UK Home Office. Transgenic Thy1-GFP-M mice (line 007788, Jackson Laboratories) expressing eGFP in sparse subsets of neurons were used for *in vivo* imaging of dendritic spines. Animals aged 4–6 weeks were premedicated with intraperitoneal injections of dexamethasone (Dexadreson, 4 μg), atropine (Atrocare, 1 μg) and carprofen (Rimadyl, 0.15 μg). General anesthesia was induced by an intraperitoneal injection of fentanyl (Sublimaze, 0.05 mg/kg), midazolam (Hypnovel, 5 mg/kg), and medetomidine (Domitor, 0.5 mg/kg). Mice were then placed in a stereotaxic frame equipped with mouth and ear bars. Depth of anesthesia was monitored by pinching the rear foot and by observation of the respiratory pattern. Body temperature was closely monitored throughout the procedure, and kept constant at 37°C by the use of a heating mat and a temperature controller in conjunction with a rectal temperature probe. Both eyes were covered with eye ointment (Maxitrol, Alcon) to prevent corneal desiccation during the experiment. The skin over the craniotomy site was shaved and an incision was made to expose the skull, after which a small hole of 0.5 mm diameter was drilled (Foredom K.1070, Blackstone Industries, CT, USA) into the skull with a 0.4 mm drill bit. The craniotomy was centred at 1.3 mm anterior and 1.0 mm lateral to bregma. Cyanoacrylate glue (Pattex Classic, Henkel, Germany) was applied to the surrounding skull, muscle, and wound margins to prevent further bleeding. A small metal bar was attached to the skull over the left hemisphere with dental cement, which was also used to cover all exposed areas of skull. The mouse was then placed on a custom-made stage, its head fixed to the stage using the steel bar, for imaging. Subsequently, the multimode optic fibre (MMF, total diameter 125 μm) was gradually lowered up to 1.8 mm into the brain tissue targeting the dorsal striatum.

For dynamic imaging, C57BL/6 mice were injected with very small amounts (<5 nl) of a 1:1 mixture of highly diluted (1:50000-100000 in PBS) AAV1.hSyn.Cre.WPRE.hGH (Penn Vector Core) and AAV1.CAG.Flex.GCaMP6m.WPRE.bGH (Penn Vector Core) into the right medial geniculate body of the thalamus. The thalamic stereotaxic coordinates were 2.9 mm posterior to bregma, 2.05 mm to the right of the midline and 3.0 mm from the cortical surface. Fibre-imaging was performed 3-4 weeks after GCaMP6m viral injections. The pre-imaging surgery was as described above, but with a different anaesthetic regime: general anesthesia was induced with ketamine (100 mg/kg, Vetalar) and medetomidine (140 μmg/kg), and ketamine (50 mg/kg/h) and medetomidine (0.07 mg/kg/h) were regularly topped up at 30 min intervals to maintain a stable level of anaesthesia throughout the experiment. Sound presentation was performed through a free-field loudspeaker (Tucker-Davis Technologies) placed near the ear canal of the mouse’s left ear. Stimuli were single 16kHz pure tones of 100 milliseconds duration.

At the end of the imaging session, the mouse was given an overdose of sodium pentobarbital (240 mg/kg) prior to transcardial perfusion with phosphate buffered saline (PBS) and then 4 % paraformaldehyde. The fixed brain was then extracted and section to confirm the location of the fibre tract.

## Acknowledgements

MP and TC acknowledge support from the University of Dundee and Scottish Universities Physics Alliance (PaLS initiative). TC acknowledges support from the European Regional Development Fund, Project No. CZ.02.1.01/0.0/0.0/15 003/0000476. SAVL, VK, ZP and NE acknowledge support from the John Fell Fund, the BBSRC (TDRF) and the MRC (UK). We thank V. De Paola (Imperial College) for providing the transgenic Thy1-eGFP-M mouse line.

## Author contributions

SAVL, VK, ZP and NE conducted all experiments. MP and TC designed and constructed the experimental geometry and developed the controlling interface. All authors contributed to writing the manuscript and analysis of the data. TC and NE led the project.

