## Supplementary Materials for "Minimally invasive deep-brain imaging through a 50 μm-core multimode fibre"

**Supplementary Table 1.** List of studies, included as data points in Figure 1b, which implement various forms of *in vivo* light-based brain imaging.

| # | Reference |
|---|---|
| | **Fibre bundle** |
| 2 | Goto, A., Nakahara, I., Yamaguchi, T., Kamioka, Y., Sumiyama, K., Matsuda, M., ... & Funabiki, K. (2015). Circuit-dependent striatal PKA and ERK signaling underlies rapid behavioral shift in mating reaction of male mice. *Proceedings of the National Academy of Sciences*, *112*(21), 6718-6723. |
| 3 | Yamaguchi, T., Goto, A., Nakahara, I., Yawata, S., Hikida, T., Matsuda, M., ... & Nakanishi, S. (2015). Role of PKA signaling in D2 receptor-expressing neurons in the core of the nucleus accumbens in aversive learning. *Proceedings of the National Academy of Sciences*, *112*(36), 11383-11388. |
| 4 | Chen, X., Cao, H., Saraf, A., Zweifel, L. S., & Storm, D. R. (2015). Overexpression of the type 1 adenylyl cyclase in the forebrain leads to deficits of behavioral inhibition. *Journal of Neuroscience*, *35*(1), 339-351. |
| 5 | Soden, M. E., Jones, G. L., Sanford, C. A., Chung, A. S., Güler, A. D., Chavkin, C., ... & Zweifel, L. S. (2013). Disruption of dopamine neuron activity pattern regulation through selective expression of a human KCNN3 mutation. *Neuron*, *80*(4), 997-1009. |
| 6 | Szabo, V., Ventalon, C., De Sars, V., Bradley, J., & Emiliani, V. (2014). Spatially selective holographic photoactivation and functional fluorescence imaging in freely behaving mice with a fiberscope. *Neuron*, *84*(6), 1157-1169. |
| 7 | Vincent, P., Maskos, U., Charvet, I., Bourgeais, L., Stoppini, L., Leresche, N., ... & Paupardin-Tritsch, D. (2006). Live imaging of neural structure and function by fibred fluorescence microscopy. *EMBO reports*, *7*(11), 1154-1161. |
| 8 | Bharali, D. J., Klejbor, I., Stachowiak, E. K., Dutta, P., Roy, I., Kaur, N., ... & Stachowiak, M. K. (2005). Organically modified silica nanoparticles: a nonviral vector for in vivo gene delivery and expression in the brain. *Proceedings of the National Academy of Sciences of the United States of America*, *102*(32), 11539-11544. |
| | **GRIN lens** |
| 1 | Murayama, M., Pérez-Garci, E., Nevian, T., Bock, T., Senn, W., & Larkum, M. E. (2009). Dendritic encoding of sensory stimuli controlled by deep cortical interneurons. *Nature*, *457*(7233), 1137. |
| 9 | Cox, J., Pinto, L., & Dan, Y. (2016). Calcium imaging of sleep–wake related neuronal activity in the dorsal pons. *Nature communications*, *7*, 10763. |
| 10 | Betley, J. N., Xu, S., Cao, Z. F. H., Gong, R., Magnus, C. J., Yu, Y., & Sternson, S. M. (2015). Neurons for hunger and thirst transmit a negative-valence teaching signal. *Nature*, *521*(7551), 180. |
| 11 | Kitamura, T., Sun, C., Martin, J., Kitch, L. J., Schnitzer, M. J., & Tonegawa, S. (2015). Entorhinal cortical ocean cells encode specific contexts and drive context-specific fear memory. *Neuron*, *87*(6), 1317-1331. |

| | |
|---|---|
| 12 | Sun, C., Kitamura, T., Yamamoto, J., Martin, J., Pignatelli, M., Kitch, L. J., ... & Tonegawa, S. (2015). Distinct speed dependence of entorhinal island and ocean cells, including respective grid cells. *Proceedings of the National Academy of Sciences*, *112*(30), 9466-9471. |
| 13 | Flusberg, B. A., Nimmerjahn, A., Cocker, E. D., Mukamel, E. A., Barretto, R. P., Ko, T. H., ... & Schnitzer, M. J. (2008). High-speed, miniaturized fluorescence microscopy in freely moving mice. *Nature methods*, *5*(11), 935. |
| 14 | Ghosh, K. K., Burns, L. D., Cocker, E. D., Nimmerjahn, A., Ziv, Y., El Gamal, A., & Schnitzer, M. J. (2011). Miniaturized integration of a fluorescence microscope. *Nature methods*, *8*(10), 871. |
| 15 | Ziv, Y., Burns, L. D., Cocker, E. D., Hamel, E. O., Ghosh, K. K., Kitch, L. J., ... & Schnitzer, M. J. (2013). Long-term dynamics of CA1 hippocampal place codes. *Nature neuroscience*, *16*(3), 264. |
| 16 | Jennings, J. H., Ung, R. L., Resendez, S. L., Stamatakis, A. M., Taylor, J. G., Huang, J., ... & Ramakrishnan, C. (2015). Visualizing hypothalamic network dynamics for appetitive and consummatory behaviors. *Cell*, *160*(3), 516-527. |
| 18 | Pinto, L., & Dan, Y. (2015). Cell-type-specific activity in prefrontal cortex during goal-directed behavior. *Neuron*, *87*(2), 437-450. |
| 23 | Barretto, R. P., Messerschmidt, B., & Schnitzer, M. J. (2009). In vivo fluorescence imaging with high-resolution microlenses. *Nature methods*, *6*(7), 511. |
| 17, 24 | Barretto, R. P., Ko, T. H., Jung, J. C., Wang, T. J., Capps, G., Waters, A. C., ... & Schnitzer, M. J. (2011). Time-lapse imaging of disease progression in deep brain areas using fluorescence microendoscopy. *Nature medicine*, *17*(2), 223. |
| 25, 26 | Attardo, A., Fitzgerald, J. E., & Schnitzer, M. J. (2015). Impermanence of dendritic spines in live adult CA1 hippocampus. *Nature*, *523*(7562), 592. |
| | **Objective** |
| 19 | Villette, V., Malvache, A., Tressard, T., Dupuy, N., & Cossart, R. (2015). Internally recurring hippocampal sequences as a population template of spatiotemporal information. *Neuron*, *88*(2), 357-366. |
| 20 | Danielson, N. B., Kaifosh, P., Zaremba, J. D., Lovett-Barron, M., Tsai, J., Denny, C. A., ... & Losonczy, A. (2016). Distinct contribution of adult-born hippocampal granule cells to context encoding. *Neuron*, *90*(1), 101-112. |
| 21 | Dombeck, D. A., Harvey, C. D., Tian, L., Looger, L. L., & Tank, D. W. (2010). Functional imaging of hippocampal place cells at cellular resolution during virtual navigation. *Nature neuroscience*, *13*(11), 1433. |
| 22 | Busche, M. A., Chen, X., Henning, H. A., Reichwald, J., Staufenbiel, M., Sakmann, B., & Konnerth, A. (2012). Critical role of soluble amyloid-β for early hippocampal hyperactivity in a mouse model of Alzheimer’s disease. *Proceedings of the National Academy of Sciences*, *109*(22), 8740-8745. |
| 27 | Mizrahi, A., Crowley, J. C., Shtoyerman, E., & Katz, L. C. (2004). High-resolution in vivo imaging of hippocampal dendrites and spines. *Journal of Neuroscience*, *24*(13), 3147-3151. |
| 28 | Rickgauer, J. P., Deisseroth, K., & Tank, D. W. (2014). Simultaneous cellular-resolution optical perturbation and imaging of place cell firing fields. *Nature neuroscience*, *17*(12), 1816. |

| | |
|---|---|
| 29 | Rajasethupathy, P., Sankaran, S., Marshel, J. H., Kim, C. K., Ferenczi, E., Lee, S. Y., ... & Liston, C. (2015). Projections from neocortex mediate top-down control of memory retrieval. *Nature*, *526*(7575), 653. |
| 30 | Lovett-Barron, M., Kaifosh, P., Kheirbek, M. A., Danielson, N., Zaremba, J. D., Reardon, T. R., ... & Losonczy, A. (2014). Dendritic inhibition in the hippocampus supports fear learning. *Science*, *343*(6173), 857-863. |
| 31 | Sheffield, M. E., & Dombeck, D. A. (2015). Calcium transient prevalence across the dendritic arbour predicts place field properties. *Nature*, *517*(7533), 200. |
| 32 | Gu, L., Kleiber, S., Schmid, L., Nebeling, F., Chamoun, M., Steffen, J., ... & Fuhrmann, M. (2014). Long-term in vivo imaging of dendritic spines in the hippocampus reveals structural plasticity. *Journal of Neuroscience*, *34*(42), 13948-13953. |
| 33 | Kaifosh, P., Lovett-Barron, M., Turi, G. F., Reardon, T. R., & Losonczy, A. (2013). Septo-hippocampal GABAergic signaling across multiple modalities in awake mice. *Nature neuroscience*, *16*(9), 1182. |
